# Analysis of glutamine synthetase target-site mutations and their role in endowing glufosinate-ammonium resistance

**DOI:** 10.1101/2025.11.01.685562

**Authors:** Aimone Porri, Susee Sudhakar, Matheus M Noguera, Michael Betz, Jens Lerchl, Franck E Dayan, Jason K Norsworthy

## Abstract

Glufosinate-ammonium (GFA) is a key non-selective herbicide for controlling *Amaranthus palmeri* and other weeds by targeting glutamine synthetase (GS). GS copy number, expression, sequence polymorphisms, and enzymatic properties in a GFA-resistant population (CCR) were studied. Digital PCR revealed no major GS amplification or target upregulation: most CCR plants had copy numbers and expression comparable to the susceptible reference, with only minor increases in GS2.1 and GS2.2 in a few individuals. Sequencing identified a non-synonymous substitution, G255D, in GS2.2 within a conserved region adjacent to the GFA-binding site. G255D retained ∼58% of wild-type activity *in vitro* assays, but was completely insensitive to GFA, with no measurable inhibition at tested concentrations. However, ectopic expression of G255D in *Arabidopsis thaliana* did not confer GFA tolerance, indicating the mutation alone is insufficient for resistance. *In vitro* analysis of the *Eleusine indica* GS1.1 S59G substitution revealed increased catalytic activity without affecting GFA sensitivity. A mutational panel of GS1.1 variants showed that substitutions at E131, E192, G245, H249, R291, R311, and R332 abolished enzyme activity or inhibitor sensitivity, with most variants retaining <2% of wild-type function. Among a broader set of predicted GS1.1 variants, high resistance indices were consistently linked to strong reductions in catalytic efficiency, underscoring the fitness costs of target-site alterations. Collectively, GS2.2 G255D appears to be a rare substitution combining substantial residual activity with complete GFA insensitivity and suggest that resistance via target-site modifications studied is constrained by trade-offs between catalytic function and herbicide binding.

## 1. Introduction

Glufosinate-ammonium (GFA) is a non-selective, postemergence herbicide widely utilized for broad-spectrum control of weeds. It targets glutamine synthetase (GS; EC 6.3.1.2), a key enzyme in nitrogen metabolism that catalyzes the ATP-dependent condensation of glutamate and ammonia to form glutamine.^1^ In higher plants, GS exists at two different locations: GS1 in the cytosol and GS2 in the chloroplast.^2^ GS2 typically dominates foliar tissues of C3 species due to its role in photorespiratory ammonia reassimilation, while both GS1 and GS2 contribute to nitrogen processing in C4 species such as Palmer amaranth (*Amaranthus palmeri* S. Watson), where GS1 activity may account for a large proportion of total GS activity.^3,4^ Inhibition of GS leads to toxic ammonia accumulation and disrupted photorespiration.^5^ Though ammonia accumulation was initially considered the primary cause of glufosinate-induced cell death, recent studies indicate that reactive oxygen species (ROS) generation and membrane lipid peroxidation contribute greatly to its phytotoxic effects.^6^

Commercialized in the 1990s, glufosinate use expanded following the introduction of transgenic crops carrying *bar* or *pat*, which encode phosphinothricin acetyltransferase to detoxify glufosinate by converting it to *N*-acetyl-glufosinate.^7^ This trait is now incorporated into multiple cropping systems, including LibertyLink®, Enlist™, and XtendFlex®. In the United States, GFA applications in soybean [*Glycine max* (L.) Merr.] increased from approximately 1.08 million kg in 2012 to 6.54 million kg in 2020, highlighting its growing role in managing herbicide-resistant weeds.^8^

Evolution of GFA resistance in weeds has been relatively limited compared to other herbicide sites of action, with only five reported GFA-resistant species: goosegrass [*Eleusine indica* (L.) Gaertn.], Italian ryegrass [*Lolium perenne* L. ssp. *multiflorum* (Lam.) Husnot], perennial ryegrass (*Lolium perenne* L.), rigid ryegrass (*Lolium rigidum* Gaudin), and *A. palmeri*.^9–16^ Among these, *A. palmeri* is the only broadleaf species with confirmed resistance. As a dioecious, highly fecund C4 weed with extensive gene flow and rapid adaptation capacity, *A. palmeri* presents a substantial threat to sustainable chemical control. Resistance to GFA in *A. palmeri* has been confirmed in populations from Arkansas, Missouri, and North Carolina, including accessions with multiple herbicide resistances.^16–19^

Mechanisms of herbicide resistance in weeds are broadly classified into target-site resistance (TSR) and non-target-site resistance (NTSR). TSR mechanisms include point mutations in the herbicide target gene, gene amplification, or overexpression of the target protein, all of which reduce herbicide binding efficacy or increase the availability of the target. NTSR mechanisms encompass lower absorption or translocation, vacuolar sequestration, or enhanced metabolic degradation, often mediated by cytochrome P450 monooxygenases (P450s), glutathione-S-transferases (GSTs), or other detoxification pathways.^20^ Glufosinate uptake can vary depending on the herbicide applied in combination, with certain mixtures reduce absorption in species such as *A. palmeri* and barnyardgrass [*Echinochloa. crus-galli* (L.) P. Beauv.].^21^ In *A. palmeri*, the most validated mechanism of glufosinate resistance to date involves GS2 overexpression and gene amplification, with limited evidence supporting a role for target-site mutations.^17,22^

A few GS mutations have been reported or engineered in relation to glufosinate sensitivity, but none have been conclusively linked to field-evolved resistance. A *GS2* Asp173Asn substitution in *L. multiflorum* was initially thought to confer resistance but was later shown to be non-functional; resistance in that population was ultimately determined to be mediated by enhanced metabolism.^23^ A S59G substitution in the GS1 isoform of *E. indica* conferred glufosinate resistance, with transgenic rice (*Oryza sativa* L.) plants expressing the mutant gene exhibiting reduced sensitivity to the herbicide.^24^ In controlled systems, substitutions such as H249Y and R295K in soybean and rice GS2 enzymes have been engineered to reduce glufosinate binding, but such mutations have not been observed in weed species under field selection.^25,26^

Recent research has shown that extrachromosomal circular DNA (eccDNA) is a novel mechanism underlying gene amplification in certain herbicide-resistant weeds, such as in glyphosate-resistant *A. palmeri*.^27^ Similarly, in a GFA-resistant *A. palmeri* population from Arkansas (MSR2), GS2.1 and GS2.2 isoforms were co-amplified on an eccDNA structure.^28^ This finding suggests that eccDNA-mediated gene amplification may serve as a flexible and rapid adaptation strategy in *A. palmeri*, allowing high-copy gene expression outside the constraints of chromosomal regulation. However, the presence of GS2-containing eccDNA has not been confirmed in all resistant populations, and resistance mechanisms are likely to be heterogeneous across regions.

The present study investigates the response of a distinct *A. palmeri* population from Arkansas (CCR) to GFA. Sequencing of the chloroplastic GS2 gene in this population revealed a glycine-to-aspartic acid substitution at position 255 (G255D), located in a conserved region of the enzyme. The G255D substitution was detected only in non-surviving plants and in heterozygous form, and it was absent in GFA survivors. No GS2 copy number variation was detected among CCR survivors. Functional enzyme assays confirmed that the G255D substitution does not confer GFA resistance.

In addition to G255D, a broader set of putative GS1 mutations was analyzed to assess their potential impact on enzyme function and herbicide interaction. This curated set provides a valuable molecular reference for future investigations. If field-evolved GFA resistance is later linked to specific GS mutations, these data may help differentiate resistance-conferring variants from neutral polymorphisms. The present study 1) evaluates GFA sensitivity in the CCR population and characterizes the G255D variant, and 2) contributes to a clearer understanding of resistance mechanisms in *A. palmeri*. As GFA continues to play a central role in weed management, the timely identification and tracking of resistance mechanisms remain essential for preserving its long-term utility.

## 2. Material and methods

### 2.1 Plant material

The *A. palmeri* CCR population and the susceptible reference were grown in a greenhouse under a 14:10 h light:dark photoperiod. Natural light was supplemented with high-pressure sodium lamps providing a photon flux density of approximately 1100 µmol m⁻² s⁻¹ during daylight hours. Day/night temperatures were maintained at 30/25 °C, respectively. Twenty-three untreated CCR plants were collected for molecular characterization of resistance.

### 2.2 Preparation of DNA

Leaf samples of 0.5 cm² each were transferred into a sample tube (Collection microtubes; Qiagen, Hilden, Germany). Subsequently, the samples were homogenized in a shaker mill (TissueLyser II; Qiagen, Hilden, Germany) with steel beads. The process of DNA extraction was carried out in a KingFisher Flex Magnetic Particle Processor (Thermo Fisher Scientific, Schwerte, Germany), employing the Chemagic Plant 400 kit (Perkin Elmer, Rodgau, Germany), in accordance with the manufacturer’s instructions (with modifications implemented by IDENTXX GmbH).

### 2.3 Preparation of total RNA

Frozen leaf samples (0.5 cm² each) were transferred into a sample tube. Subsequently, the samples were homogenized with steel beads in a shaker mill (TissueLyser II; Qiagen, Hilden, Germany). The total RNA extraction was carried out in the KingFisher Flex Magnetic Particle Processor (Thermo Fisher Scientific, Schwerte, Germany), employing the MagMAX(™) Plant RNA Isolation Kit (Applied Biosystems, Darmstadt, Germany), in accordance with the manufacturer’s instructions.

### 2.4 Preparation of cDNA

The random hexamer-primed cDNA was obtained by reverse transcription of 200 ng µL^-1^ of total RNA using the PrimeScript RT Reagent Kit with gDNA Eraser (Takara Bio Europe, St-Germain-en-Laye, France) according to the manufacturer’s instructions.

### 2.5 GS copy number and expression analysis

The digital PCR (dPCR) was performed in a final volume of 12 µL using 1.48 µL of DNA or cDNA, 0.48 µL (0.2 µM) of specific primers and 0.24 µL (0.2 µM) of probe (Table S1) (biomers.net GmbH, Ulm, Germany), 3 µL of QIAcuity HighMultiplex ProbePCR Kit (Qiagen, Hilden Germany) and 4.64 µL PCR-Grade H_2_O for the quintruple dPCR. dPCR was performed in a dPCR thermal cycler (QIAcuity One, 5plex Device, Qiagen, Hilden Germany) in a 96-well nanoplate (QIAcuity Nanoplate 8.5k 96-well) under the following conditions: 2 min at 95°C and 55 cycles of 15 s denaturation at 95°C; 55 s annealing and elongation at 60°C. The following conditions were met during the imaging of the partitions. The FAM and HEX experiments were conducted with an exposure time of 500 milliseconds, with the gain set to 6. The ROX experiment utilized an exposure time of 300 milliseconds, also with the gain set to 4. The TAMRA experiment utilized an exposure time of 400 milliseconds, also with the gain set to 6. The Cy5 experiment utilized an exposure time of 400 milliseconds, also with the gain set to 8. The assessment of copy number variation and expression analysis was conducted by employing Qiagen’s QIAcuity Software Suite, version 3.1.0.0. This process entailed the utilization of positive, negative, and no-template control wells (NTC), along with the determination of sample thresholds.

### 2.6 Identification of target-Site mutations in *A. palmeri* CCR population GS isoforms via cDNA sequencing

The amplification of the coding sequences was performed in a final volume of 25 µL using 5 µL of the random hexamer primed cDNA and 1 µL (10 pmol) of specific primers (Table S2), 12.5 of MyFi™ DNA Polymerase (Bioline GmbH, Luckenwalde, Germany) and 6.5 µL PCR-Grade H2O. DNA was amplified in a PCR thermal cycler (T100 PCR thermal cycler, Bio-Rad Laboratories GmbH, Germany) under the following conditions: 3 min at 95°C and 35 cycles of 30 s denaturation at 95°C; 30 s annealing at primer specific temperature and 2 min elongation at 72°C; and a final elongation step at 72°C for 5 min. Aliquots were taken and analysed on 1.5% agarose gels. The PCR products were sequenced from both sites via Sanger sequencing (SeqLab-Microsynth, Göttingen, Germany). Chromatograms were processed in Geneious v. 9.1.8 (Biomatters, Auckland, New Zealand), and sequences were aligned against wild-type GS coding sequences. Resulting nucleotide substitutions were translated into amino acid changes to identify potential target-site mutations

### 2.7 Protein expression and purification

The full-length of GS wild-type and variants CDS sequences form *A. palmeri* and *E. indica* were synthesized de novo and cloned into the pRSetB expression vector (Invitrogen, Carlsbad, CA, USA) using BamHI and HindIII restriction sites. An N-terminal hexahistidine tag was included to facilitate purification. Recombinant constructs were transformed into *Escherichia coli* strain BL21(DE3)pLysS (Novagen, EMD Millipore, Billerica, MA, USA), and transformants were selected on LB agar containing 100 µg mL⁻¹ ampicillin and 34 µg mL⁻¹ chloramphenicol.

For protein expression, a single colony was used to inoculate 3 mL LB medium with antibiotics and incubated at 37°C with shaking (200 rpm) for 6 h. A 20-µL aliquot of this pre-culture was transferred into 20 mL fresh LB medium and incubated overnight. The next day, 100 µL of the overnight culture was inoculated into 100 mL ZYM-5052 autoinduction medium (see Supplemental Data for preparation), supplemented with antibiotics, and incubated at 37°C for 5 h, followed by 21 h at 25°C.

Cells were harvested by centrifugation at 6,000 × g for 30 min at 4°C. Cell pellets were resuspended in PPO lysis buffer [10 mL g⁻¹ pellet; 50 mM NaH₂PO₄, 100 mM NaCl, 5 mM imidazole, 5% (v/v) glycerol, pH 7.5, 20 mg mL⁻¹ lysozyme, 30 U mL⁻¹ DNase I] supplemented with protease inhibitors (complete EDTA-free; Roche Diagnostics, Mannheim, Germany). Suspensions were sonicated on ice (3 min, 90% amplitude, 30-s intervals). Cell debris was removed by centrifugation at 38,000 × g for 30 min at 4°C, and 2 mL of 5 M NaCl was added to the clarified supernatant.

For purification, a 500-µL bed volume of HisPur Ni-NTA resin (Thermo Fisher Scientific, IL, USA) was equilibrated with buffer (20 mM NaH₂PO₄, 50 mM NaCl, 5 mM imidazole, 5 mM MgCl₂, 17% glycerol, 0.1 mM EDTA, pH 8.0). The supernatant was passed through the resin, washed with 5.6 mL wash buffer (20 mM NaH₂PO₄, 50 mM NaCl, 5 mM imidazole, 17% glycerol, pH 7.5), and the bound protein was eluted with 1 mL elution buffer (20 mM NaH₂PO₄, 50 mM NaCl, 250 mM imidazole, 17% glycerol, pH 7.5). Protein concentrations were determined using a Scandrop nanovolume spectrophotometer (Analytikjena, Life Science, Germany). Purity and solubility were assessed by SDS-PAGE (10%) using 2.5 µg protein per lane.

### 2.8 Glutamine synthetase enzyme activity

GS activity was determined spectrophotometrically following the Sigma-Aldrich enzymatic assay protocol. The reaction couples the ATP-dependent conversion of L-glutamate and ammonium to L-glutamine with pyruvate kinase (PK) and L-lactate dehydrogenase (LDH), such that oxidation of β-NADH to β-NAD⁺ can be monitored at 340 nm. Recombinant GS from *A. palmeri* was expressed in Escherichia coli, purified by affinity chromatography, and quantified by the Bradford method using bovine serum albumin as standard. Assays were conducted at 37°C in 3.00 mL total volume containing 34.1 mM imidazole (pH 7.1), 102 mM sodium L-glutamate, 8.5 mM ATP, 1.1 mM phosphoenolpyruvate, 60 mM MgCl₂, 18.9 mM KCl, 45 mM NH₄Cl, 0.25 mM β-NADH, 28 U PK, 40 U LDH, and 0.4–0.8 U recombinant GS. Reaction mixtures were equilibrated to 37°C and initiated by the addition of enzyme, and the decrease in absorbance at 340 nm was recorded for 5–10 min; only the linear portion of the curve was used for activity calculations. For inhibition assays, GFA was added directly to the reaction mixture to final concentrations ranging from 10⁻⁸ M to 10⁻² M, using 2.5-fold serial dilutions, with a no-inhibitor control included in each run. Activity was calculated using an extinction coefficient for NADH of 6.22 mM⁻¹ cm⁻¹ and expressed as units per mg protein, where one unit corresponds to the formation of 1 µmol of L-glutamine in 15 min at pH 7.1 and 37°C. Background activity was corrected by including blank assays without GS. Percent inhibition was calculated relative to untreated (positive) and no-enzyme (negative) controls. Assays were conducted in duplicate. IC₅₀ values (concentration of inhibitor reducing GS activity by 50%) were estimated by nonlinear regression (three- or four-parameter log-logistic models). Resistance factors were calculated by dividing the IC₅₀ of each mutant variant by that of wild-type GS.

### 2.9 Transgene preparation and Arabidopsis transformation

Mutant GS variants G2.2 G255D from *A. palmeri* and G254D from Arabidopsis were cloned into the RTP6557 transformation vector, which was subsequently introduced into *Agrobacterium tumefaciens* strain C58C1pMP90. The construct also carried an acetolactate synthase (ALS) herbicide-resistance gene as a selectable marker, ensuring that Arabidopsis seedlings tested for resistance to GFA herbicide expressed the transgene.

Agrobacterium cultures were initiated 1 d prior to transformation by inoculating 1 mL of glycerol stock into 250 mL of YEB medium (1 g L⁻¹ yeast extract, 5 g L⁻¹ beef extract, 5 g L⁻¹ peptone, 5 g L⁻¹ sucrose, and 0.49 g L⁻¹ MgSO₄·7H₂O) supplemented with the appropriate antibiotic. Cultures were grown for 12 h at 28°C with agitation at 150 rpm. On the following day, the culture density was adjusted to OD₆₀₀ = 1.0 in YEB medium, harvested by centrifugation (1600 g, 10 min), and resuspended in 150 mL of infiltration medium (2.2 g L⁻¹ Murashige & Skoog salts, 50 g L⁻¹ sucrose, 0.5 g L⁻¹ MES hydrate, and 10 µL L⁻¹ BAP [1 mg mL⁻¹ stock solution]). The pH of the infiltration medium was adjusted to 5.7–5.8.

Arabidopsis plants carrying immature floral buds were transformed by the floral-dip method, immersing inflorescences for 10 s in the bacterial suspension supplemented with Silwet-L77 at 0.05% vv^-1^. After dipping, plants were maintained overnight under high humidity and low light, then transferred to long-day conditions until maturity. Once siliques had yellowed, seeds were harvested into paper bags. T₁ seeds were collected, placed into Falcon tubes, and stored at 4°C.

After ≥14 d of cold storage, T₁ seeds were sown and stratified as previously described. Putative transgenic seedlings were selected by treatment with a 20 ppm imazamox (technical grade) solution. Plants were grown under short-day conditions for 12–14 d until the four-leaf stage, at which point resistant individuals were transplanted into 6 × 6 cm pots containing GS90 soil. Plants were grown for an additional 10 d, then shifted to long-day growth conditions 1 d prior to herbicide application.

Herbicide treatments were applied at the ten-leaf stage using a spray chamber calibrated to deliver 375 L ha⁻¹ of spray solution. GFA herbicide was applied at rates of 100, 500, and 1000 g active ingredient (a.i.) ha⁻¹, with DASH HC (BASF, ref. ID no. 30059102; 349 g L⁻¹ oil [fatty acid esters] and 209 g L⁻¹ alkoxylated alcohols-phosphate esters) as an adjuvant. Control plants received spray solution containing adjuvant only. Herbicide efficacy was visually evaluated 7 d after application.

### 2.10 Molecular Modeling

A homology model of AMAPA cytosolic GS1.1 was built to identify residues for targeted experiments on GFA binding, with emphasis on protein–protein interactions that can influence herbicide sensitivity. The maize GS1.1 crystal structure co-crystallized with ADP and methionine sulfoximine phosphate (PDB 2D3A served as the template.^29^ Model construction and placement of GFA into the site corresponding to methionine sulfoximine phosphate were performed in MOE using default settings. [*Molecular Operating Environment (MOE), 2024.06; Chemical Computing Group ULC, 1010 Sherbrooke St. West, Suite #910, Montreal, QC, Canada, H3A 2R7, 2024.]* For the template region resolved in the protein crystal structure (residues C3–P356), AMAPA cytosolic GS1.1 shares 88.1% sequence identity and 95.5% similarity with maize; residues directly contacting GFA are fully conserved, indicating preserved active site geometry and identical ligand interactions. However, sensitivity or resistance to GFA is also influenced by the stability and architecture of subunit interfaces in the GS oligomer. One principal interface is mainly formed by Nterminal residues (R34–E69; Table 3), and a second interface involves a central sequence segment.

To evaluate how amino acid substitutions might affect these interfaces, interfacial residues were mapped from the homology model and a multiple sequence alignment, and prioritized candidate mutations based on evolutionary conservation and single nucleotide polymorphisms (SNPs). Selected substitutions were modeled in silico and assessed by molecular modeling to estimate their effects on interface stabilization. Analyses focused on potential changes to intermolecular contacts critical for oligomer integrity — notably polar and charge assisted hydrogen bonds (e.g. E69 to R316) and hydrophobic patches (e.g. V161, A163) that could alter oligomer packing or ligand access. These structure-based considerations provide a rationale for selecting residues for experimental validation to dissect how active site binding and oligomeric interfaces jointly determine GFA sensitivity and resistance.

## 3. Results

### 3.1 Copy number variation and mRNA abundance of GS isoforms in the *A. palmeri* CCR population

Digital PCR (dPCR) analysis was employed to detect differences in gene copy number and mRNA abundance among the four GS isoforms in 23 untreated CCR plants, using an herbicide-sensitive plant as reference (fold change = 1.0). For the majority of CCR plants, copy number values for all isoforms were close to reference, generally falling between 0.8 and 1.5, indicating no substantial gene amplification (Fig. 1A). Few CCR plants exhibited slight deviations. CCR13, CCR17, and CCR22 showed a slight increase in GS2.2 CN, with fold changes reaching upto 2.0-fold. In CCR13 and CCR22 plants, a mild elevation in copies was observed also for GS2.1, suggesting that amplification in these isoforms is not widespread but can occur in certain CCR individuals. No plant demonstrated extreme amplification (>10-fold) for any GS isoform.

**FIGURE 1.**
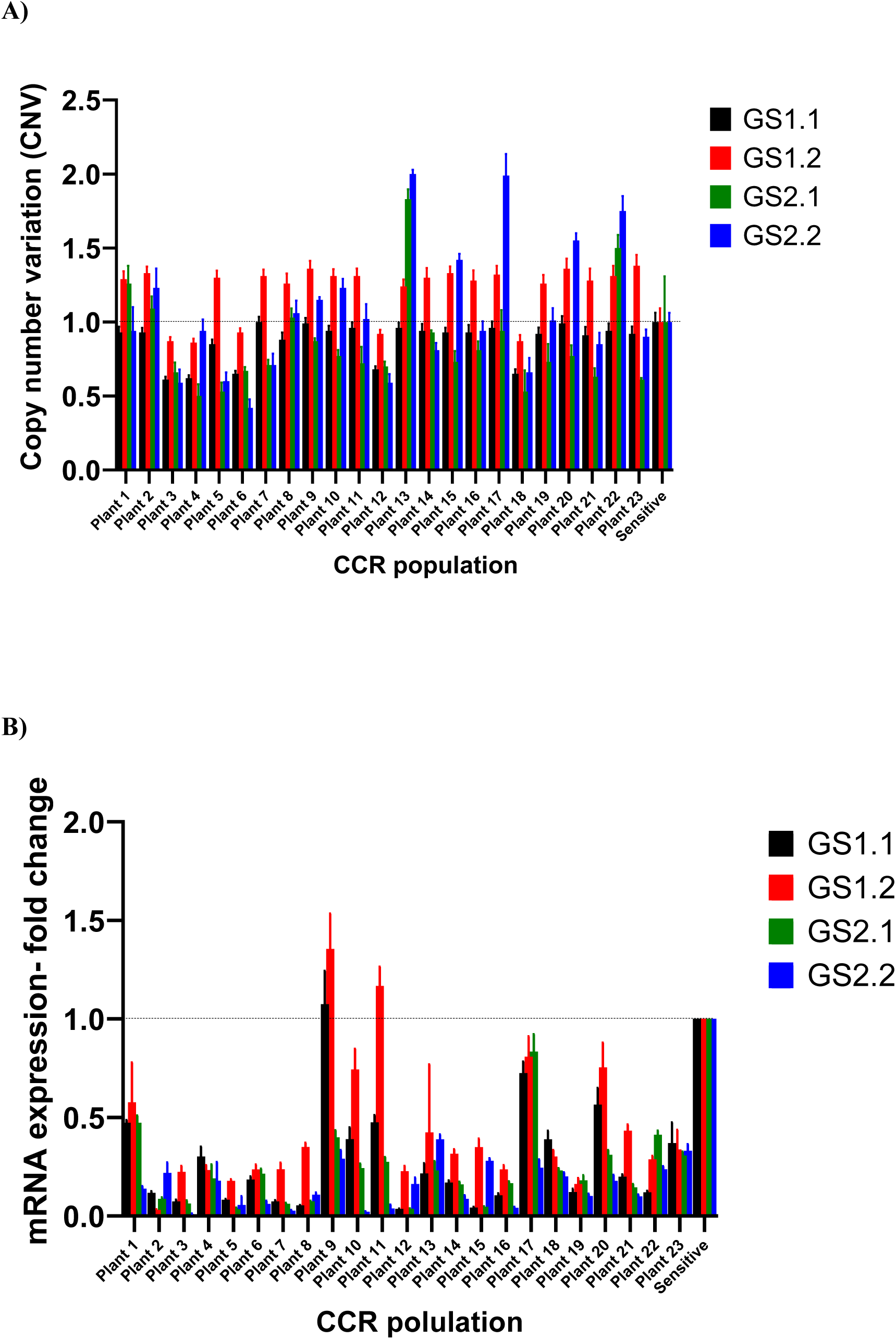
Copy number variation (CNV) (A) and mRNA abundance (B) of *Amaranthus palmeri* GS in CCR plants determined by digital PCR (dPCR). CNV values and expression fold change are shown for GS1.1 (black), GS1.2 (red), GS2.1 (green), and GS2.2 (blue). Values were calculated using the ΔΔCₜ method with a known glufosinate ammonium (GFA)-sensitive *A. palmeri* sensitive plant as reference which contains only one copy of each GS isoforms. Error bars indicate confidence interval 95% (CI 95).

Expression analysis also did not detect major target upregulation with most of the plants expressing the GS isoforms similar or slightly lower to the sensitive plant (Fig. 1B). Taken together, the vast majority of CCR plants maintain GS copy numbers or expression similar to the sensitive control, with increases in GS2.1 and GS2.2 CN being found in a few individuals.

### 3.2 GS cDNA sequencing in CCR plants

The sequencing of the GS2.2 isoform in the 23 CCR plants revealed a non-synonymous substitution at position 255, where glycine (G) in the susceptible reference sequence was replaced by aspartic acid (D). This G255D substitution was present in heterozygous form in several plants (Table 1). This residue lies within a highly conserved domain of GS 2.2, near E192 (in GS1.1) or E251 (in GS2.1) (Fig. S1), a residue involved in GFA binding, suggesting that the amino acid substitution may have functional significance in inhibitor and/or substrate binding.^29^ The mutation was detected in heterozygosis in multiple CCR individuals, while others retained the wild-type glycine. Sequencing of the remaining three GS isoforms (GS1.1, GS1.2 and GS2.1) did not reveal any amino acid substitutions compared to the susceptible reference (Table 1). Given the conservation of the G255 site across GS isoforms, and the proximity with E251, the G255D substitution may alter enzyme properties and affect GFA binding to the target enzyme.

**Table 1.** GS isoforms cDNA sequencing in CCR population. Sanger sequencing was employed to detect sequence polymorphisms in 23 untreated CCR plants in all four GS isoforms. NP= No polymorphisms detected and G255D+/- indicates that the substitution was heterozygous. G255 indicates that the plant retained the wild-type allele.

| CCR<br>population | GS1.1 | GS1.2 | GS2.1 | GS2.2 |
| --- | --- | --- | --- | --- |
| Plant 1 | NP | NP | NP | G255D+/- |
| Plant 2 | NP | NP | NP | G255 |
| Plant 3 | NP | NP | NP | G255 |
| Plant 4 | NP | NP | NP | G255D+/- |
| Plant 5 | NP | NP | NP | G255D+/- |
| Plant 6 | NP | NP | NP | G255D+/- |
| Plant 7 | NP | NP | NP | G255D+/- |
| Plant 8 | NP | NP | NP | G255D+/- |
| Plant 9 | NP | NP | NP | G255D+/- |
| Plant 10 | NP | NP | NP | G255D+/- |
| Plant 11 | NP | NP | NP | G255D+/- |
| Plant 12 | NP | NP | NP | G255D+/- |
| Plant 13 | NP | NP | NP | G255D+/- |
| Plant 14 | NP | NP | NP | G255D+/- |
| Plant 15 | NP | NP | NP | G255D+/- |
| Plant 16 | NP | NP | NP | G255D+/- |
| Plant 17 | NP | NP | NP | G255D+/- |
| Plant 18 | NP | NP | NP | G255 |
| Plant 19 | NP | NP | NP | G255 |
| Plant 20 | NP | NP | NP | G255 |
| Plant 21 | NP | NP | NP | G255D+/- |
| Plant 22 | NP | NP | NP | G255D+/- |
| Plant 23 | NP | NP | NP | G255D+/- |

### 3.3 Enzyme activity and inhibition profile of GS2.2 G255D

Enzyme assays comparing the wild-type GS 2.2 and the G255D variant revealed a substantial reduction in catalytic activity associated with the amino acid substitution. The wild-type GS 2.2 displayed full activity, measured at approximately −30 mOD min⁻¹ (100% activity) (Fig. 2). In contrast, the G255D variant retained only 58% of wild-type activity, corresponding to approximately −17 mOD min⁻¹. This decrease in activity indicates that the G255 residue plays a role in substrate interaction and/or catalytic efficiency. The data supports the hypothesis that the G255D substitution, located in a highly conserved domain, alters GS2.2 enzyme function. In addition, enzyme inhibition assays demonstrated a striking difference in sensitivity to GFA between the wild-type GS2.2 and the G255D variant (Fig. 3). The wild-type enzyme displayed a typical dose–response curve, with activity decreasing as GFA concentration increased, resulting in an IC₅₀ of 1.1 × 10⁻⁴ M. In contrast, the GS2.2 G255D variant maintained stable activity across the full range of GFA concentrations tested, with no measurable inhibition and an IC₅₀ value that could not be determined. This absence of inhibition indicates that the G255D substitution confers a high level of resistance to GFA *in vitro*, consistent with the residue predicted role in substrate binding and inhibitor interaction.

**FIGURE 2.**
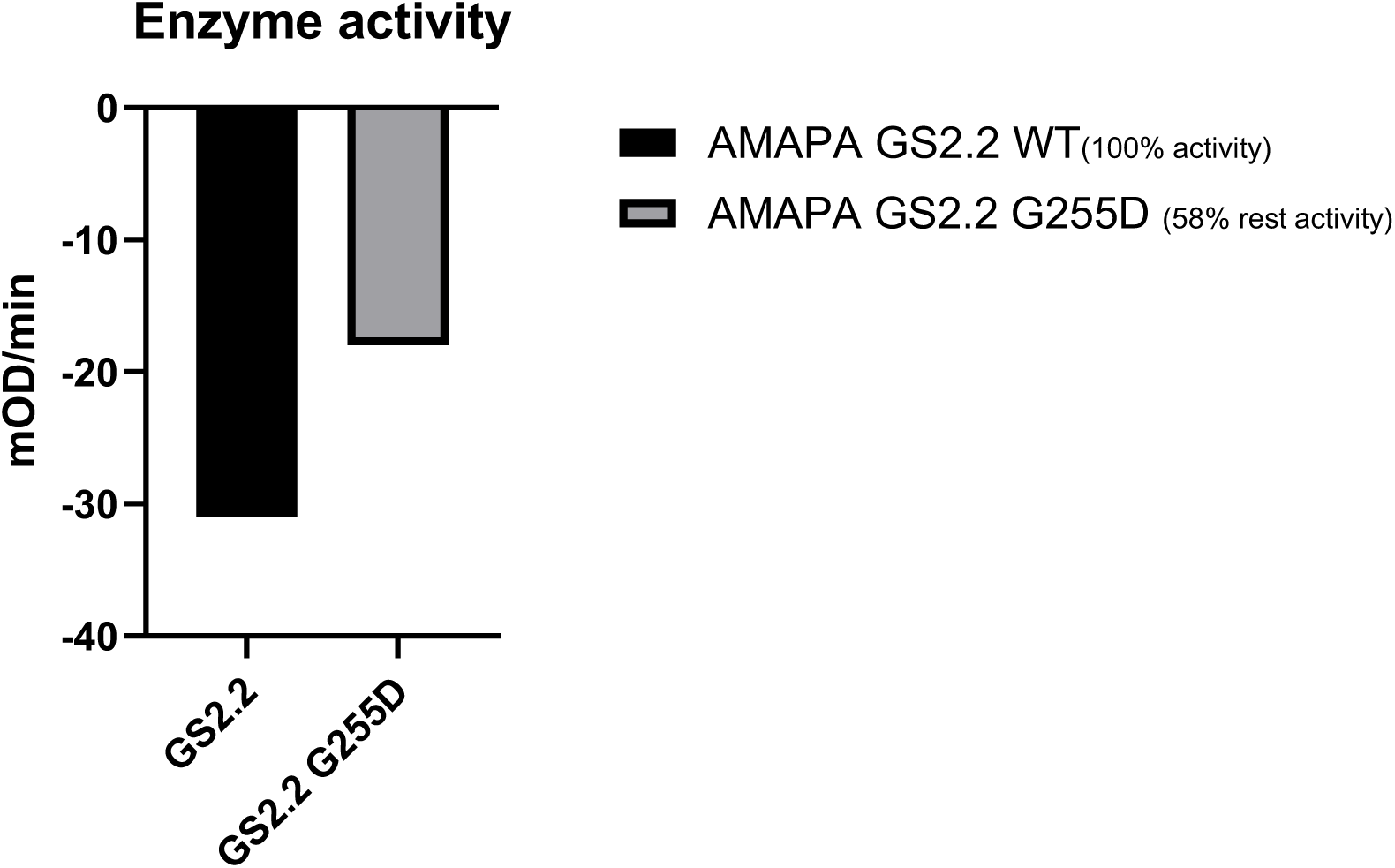
Enzymatic activity of wild-type GS2.2 and the G255D mutant. Activities were measured *in vitro* and are expressed as the change in absorbance per min (mOD min⁻¹). Wild-type GS2.2 activity was set to 100% (black bar), while the G255D variant retained 58% of WT activity (grey bar).

**FIGURE 3.**
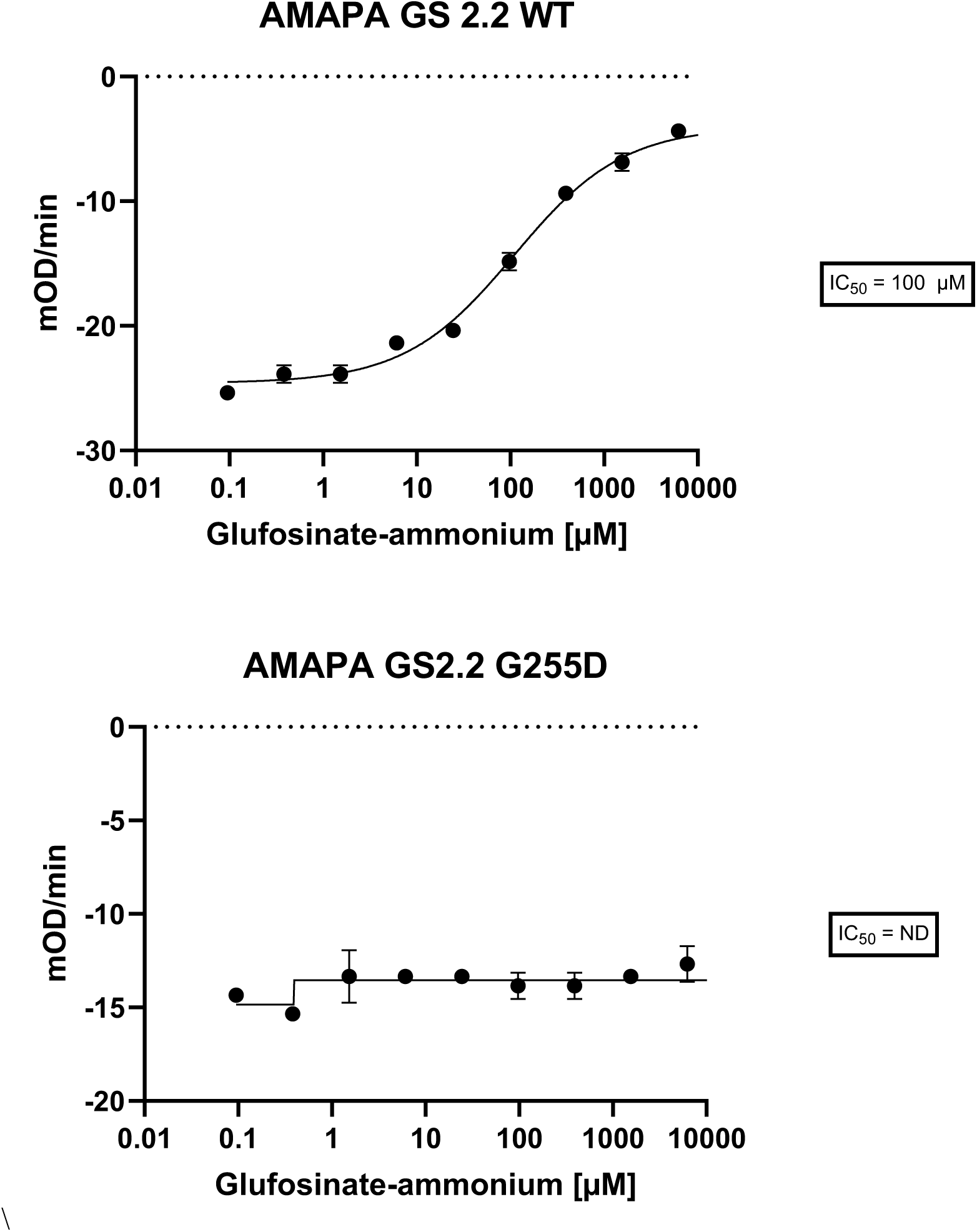
Inhibition of *Amaranthus palmeri* GS2.1 wild-type (WT) and G255D mutant enzymes by glufosinate-ammonium (GFA). Enzyme activity was measured as the change in absorbance per min (mOD min⁻¹) across a range of GFA concentrations. The WT GS2.2 enzyme (upper panel) exhibited a sigmoidal dose–response curve with an IC₅₀ of 100 µM. The G255D mutant (lower panel) showed no detectable inhibition within the tested concentration range, and an IC₅₀ could not be determined (ND). Data points represent the mean ± standard error of three replicate measurements.

### 3.4 Ectopic expression of G255D in Arabidopsis transgenics

Transgenic plants ectopically expressing the GS2.2 G255D variant were evaluated for tolerance to GFA at three application rates (100, 500, and 1000 g ha^-1^). At all concentrations tested, plants expressing the G255D variant exhibited visible injury symptoms similar to those observed in wild-type controls, including chlorosis and necrosis of young leaves (Fig. 4A). The G255D-expressing lines exhibited no improvement in plant survival or injury level compared to wild-type plants at any dose. Despite conferring strong resistance to inhibition by GFA *in vitro*, the G255D substitution alone is insufficient to provide GFA tolerance *in planta* when expressed ectopically. In addition, the G255D was also ectopically expressed using the *Arabidopsis* GS2.2 backbone (G225D of *A. palmeri* corresponds to G254D in *A. Thaliana*). Like G255D, G254D failed to confer any visual tolerance in the transgenic plants (Fig. 4B).

**FIGURE 4.**
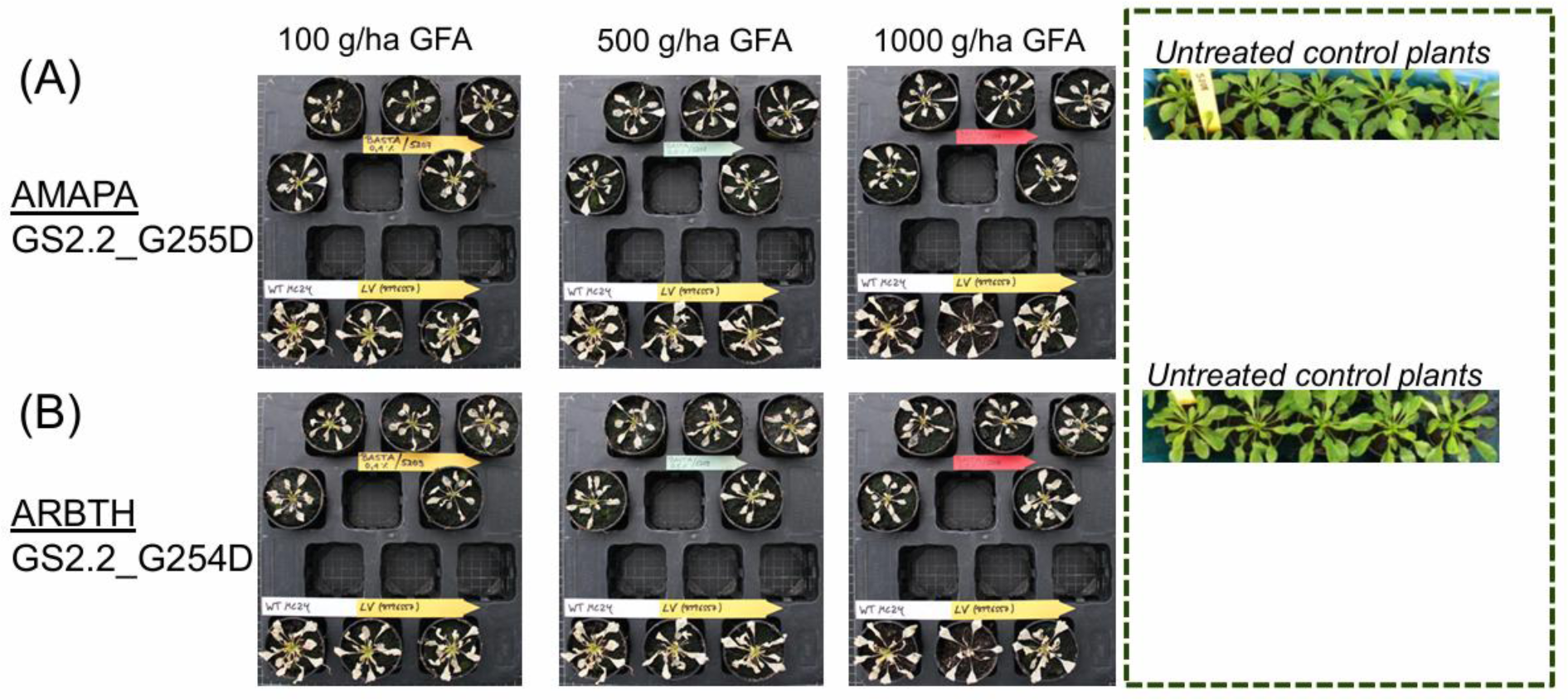
Response of (A) *Arabidopsis thaliana* plants expressing *Amaranthus palmeri* GS2.1 G255D or G254D to postemergence glufosinate-ammonium (GFA) application. Independent T₁ transgenic events expressing GS2.1_G255D A or G254D (B) (upper five pots in each tray) were sprayed at the rosette stage with GFA at 100, 500, or 1000 g a.i. ha⁻¹. Wild-type plants (WT MC24) and empty vector controls (LV RTP6557) are shown in the bottom row of each tray. Photographs were taken 10 days after treatment. Transgenic plants were confirmed by PCR detection of the AHAS resistance marker gene. Untreated control plants are also shown.

### 3.5 GS1 target-site mutations effect on GFA sensitivity

A set of cytosolic GS1.1 variants from *A. palmeri* was assayed for sensitivity to GFA. These variants were generated on residues important for GFA binding^29^ (Fig S1). Variants were created by a single nucleotide change in the codon encoding for each selected residue (Table S3). The wild-type enzyme (WT) was strongly inhibited by GFA, with an IC₅₀ of 22.5 µM and complete inhibition (100%) at 10 mM inhibitor, consistent with high susceptibility.

In contrast, many GS1.1 variants harboring substitutions at residues E131, E192, G245, H249, R291, R311, or R332 displayed extreme loss of sensitivity, with IC₅₀ values exceeding 10 mM. At saturating inhibitor concentrations, inhibition rarely exceeded 10%, and residual activities were typically <2% of WT (Table 2). Several variants (e.g. E192G, G245N/R/V, H249Q/P/Y, R311G, R332C/H/K) were catalytically inactive, precluding IC₅₀ estimation. The highest remaining activity observed was for the G245C variant (1.8%), while all the others were < 1% relative to the wild-type enzyme.

**Table 2.** Enzymatic activity and glufosinate-ammonium sensitivity of cytosolic GS1.1 variants from *Amaranthus palmeri*. Mutations were introduced into the cytosolic GS1.1 isoform and purified enzymes were assayed in vitro. IC₅₀ values represent the inhibitor concentration required to reduce activity by 50%, calculated from dose–response curves. “Inhibition at 10⁻² M” indicates the percentage inhibition at saturating glufosinate-ammonium relative to uninhibited control. “Remaining activity” represents the baseline enzymatic activity in the absence of inhibitor, expressed as a percentage of wild-type (WT) activity. Variants designated “inactive” exhibited no detectable activity under assay conditions and IC₅₀ values are not reported.

| GS1.1<br>Isoform | Mutation | Glufosinate-ammonium<br>IC<sub>50</sub> [μM]<br>P<0.0001 | Glufosinate-ammonium<br>% Inhibition at<br>saturation 10000 μM | Remaining<br>activity<br>[%] |
| --- | --- | --- | --- | --- |
| Cytosolic | WT | 22.5 | 100 | 100 |
| Cytosolic | E131A | >10000 | 8.5 | 0.04 |
| Cytosolic | E131D | >10000 | 48 | 0.2 |
| Cytosolic | E131G | >10000 | 7.2 | 0.04 |
| Cytosolic | E131K | >10000 | 0 | 0.04 |
| Cytosolic | E131Q | >10000 | 6.5 | 0.04 |
| Cytosolic | E131V | >10000 | 6.6 | 0.03 |
| Cytosolic | E192A | Enzyme inactive | - | - |
| Cytosolic | E192D | >10000 | 0 | 0.89 |
| Cytosolic | E192G | Enzyme inactive | - | - |
| Cytosolic | E192K | >10000 | 1.3 | 0.03 |
| Cytosolic | E192Q | >10000 | 0 | 0.03 |
| Cytosolic | E192V | Enzyme inactive | - | - |
| Cytosolic | G245A | Enzyme inactive | - | - |
| Cytosolic | G245C | >10000 | 0 | 1.81 |
| Cytosolic | G245S | >10000 | 9 | 0.2 |
| Cytosolic | G245N | Enzyme inactive | - | 0 |
| Cytosolic | G245R | Enzyme inactive | - | 0 |
| **Cytosolic** | G245V | Enzyme inactive | - | 0 |
| **Cytosolic** | H249L | >10000 | 7.8 | 0.03 |
| **Cytosolic** | H249N | >10000 | 17 | 0.06 |
| **Cytosolic** | H249Q | Enzyme inactive | - | - |
| **Cytosolic** | H249P | Enzyme inactive | - | - |
| **Cytosolic** | H249Y | Enzyme inactive | - | - |
| **Cytosolic** | H249R | >10000 | 29 | 0.02 |
| **Cytosolic** | R291C | >10000 | 0 | 0.07 |
| **Cytosolic** | R291S | >10000 | 0.3 | 0.08 |
| **Cytosolic** | R291G | >10000 | 0 | 0.06 |
| **Cytosolic** | R291H | >10000 | 0 | 0.1 |
| **Cytosolic** | R291L | >10000 | 0 | 0.05 |
| **Cytosolic** | R291P | >10000 | 0 | 0.04 |
| **Cytosolic** | R311G | Enzyme inactive | - | - |
| **Cytosolic** | R311L | >10000 | 14 | 0.35 |
| **Cytosolic** | R311P | >10000 | 0 | 0.16 |
| **Cytosolic** | R311Q | >10000 | 1.9 | 0.46 |
| **Cytosolic** | R332G | Enzyme inactive | - | - |
| **Cytosolic** | R332S | Enzyme inactive | - | - |
| **Cytosolic** | R332K | Enzyme inactive | - | - |
| **Cytosolic** | R332M | Enzyme inactive | - | - |
| **Cytosolic** | R332T | Enzyme inactive | - | - |
| **Cytosolic** | R332W | Enzyme inactive | - | - |

Overall, several specific mutations in cytosolic GS1.1 catalytic domain dramatically abolish sensitivity to GFA but impair catalytic efficiency simultaneously. This indicates that resistance-conferring substitutions come at a substantial cost to enzyme functionality.

A panel of additional GS1.1 variants carrying putative target-site resistance mutations in the GS1.1 subunits interaction domain, where GFA is predicted to bind, were obtained by modeling simulation (see material and methods) (Fig S2) and evaluated for both baseline catalytic activity (expressed as % of wild-type GS1.1) and resistance index (RI) to GFA (Fig. 6 and Table 3). Around 10 of the tested variants clustered near the origin, having low RIs (<5) and reduced activity, often below 40% of the wild-type level, indicating that these substitutions were either functionally neutral in terms of resistance or detrimental to enzyme performance. A smaller subset displayed moderate resistance (RI 8–12) but at the cost of substantially reduced catalytic activity (<40%). Four variants had very high GS activity ranging from 200 to 800 % of the wild-type enzyme but also low RI. Finally, another ten variants display mild and high RI (7 to 750) but severe reduction in enzyme activity below 20%. To further explore the relevance of previously reported target-site alterations, the *Eleusine indica* GS1.1 S59G mutation was also evaluated in this study for its effects on catalytic efficiency and inhibitor sensitivity.

**Table 3.** Additional mutations introduced into GS1.1 isoform of *Amaranthus palmeri*, obtained by modeling prediction. IC₅₀ values represent the inhibitor concentration required to reduce activity by 50%, calculated from dose–response curves. Remaining activity represents the baseline enzymatic activity in the absence of inhibitor, expressed as a percentage of wild-type (WT) activity.

| **GS1.1**<br>**Isoform** | **Mutation** | **Remaining**<br>**activity**<br>**[%]** | **Glufosinate-**<br>**ammonium**<br>**IC50[μM]**<br>**P<0.0001** | **Resistance**<br>**index (RI)** |
| --- | --- | --- | --- | --- |
| **Cytosolic** | **WT** | **100** | **21.7** | **1** |
| Cytosolic | R34A | 10 | 182 | 8.4 |
| Cytosolic | K36A | 14 | 248 | 11.5 |
| Cytosolic | R38E | 166 | 4.8 | 0.2 |
| Cytosolic | Y55L | 37 | 17 | 0.8 |
| Cytosolic | D56C | 5 | 3.9 | 0.2 |
| Cytosolic | D56S | 2 | 56 | 2.6 |
| Cytosolic | S58M | 105 | 94 | 4.3 |
| Cytosolic | S59N | 4 | 39.5 | 1.8 |
| Cytosolic | T60Q | 5 | 366 | 16.9 |
| Cytosolic | E69L | 39 | 246 | 11.4 |
| Cytosolic | G155F | 31 | 254 | 11.7 |
| Cytosolic | Y158A | 5 | 1401 | 64.6 |
| Cytosolic | Y158T | 29 | 76 | 3.5 |
| Cytosolic | C159H | 1 | 4046 | 186.5 |
| Cytosolic | G160T | 13 | 405 | 18.7 |
| Cytosolic | V161N | 6 | 64 | 3.0 |
| Cytosolic | V161S | 23 | 39 | 1.8 |
| Cytosolic | G162N | 2 | 489 | 22.6 |
| Cytosolic | A163Y | 101 | 54 | 2.5 |
| Cytosolic | K165S | 71 | 7.6 | 0.3 |
| Cytosolic | K165V | 209 | 33 | 1.5 |
| Cytosolic | S174E | 529 | 1 | 0.0 |
| Cytosolic | K177L | 823 | 1.3 | 0.1 |
| Cytosolic | V193T | 85 | 69 | 3.2 |
| Cytosolic | G245P | 22 | 247 | 11.4 |
| Cytosolic | E297A | 9 | 6.9 | 0.3 |
| Cytosolic | E297Q | 13 | 0.6 | 0.0 |
| Cytosolic | E297S | 19 | 0 | 0.0 |
| Cytosolic | A309L | 34 | 204 | 9.4 |
| Cytosolic | A309Y | 4 | 17355 | 799.7 |
| Cytosolic | R316A | 6 | 313 | 14.4 |
| Cytosolic | R316E | 2 | 197 | 9.1 |

### 3.6 The S59G target site mutation does not affect GFA inhibition potency

To assess whether the S59G amino acid substitution in ELEIN GS1.1 affects sensitivity to the GFA, as previously published^24^, inhibition kinetics were compared between the wild-type and mutant enzymes. Both variants displayed highly similar dose–response profiles, with enzymatic activity progressively declining as GFA concentration increased. The IC₅₀ of GFA on wild-type and S59G variant enzymes were 13μM and 19μM, respectively, representing only a minor, biologically insignificant difference with regards to the resistance mechanism (Fig. 5). Maximal inhibition was achieved at similar concentrations in both cases, and the shape of the inhibition curves was comparable, indicating no detectable change in inhibitor binding or susceptibility. However, the S59G substitution results in a higher enzyme activity compared to the wild-type enzyme (Fig S3), suggesting that the S59G substitution does not alter GS1.1 function with respect to GFA sensitivity, but it increases the overall catalytic activity of the enzyme.

**FIGURE 5.**
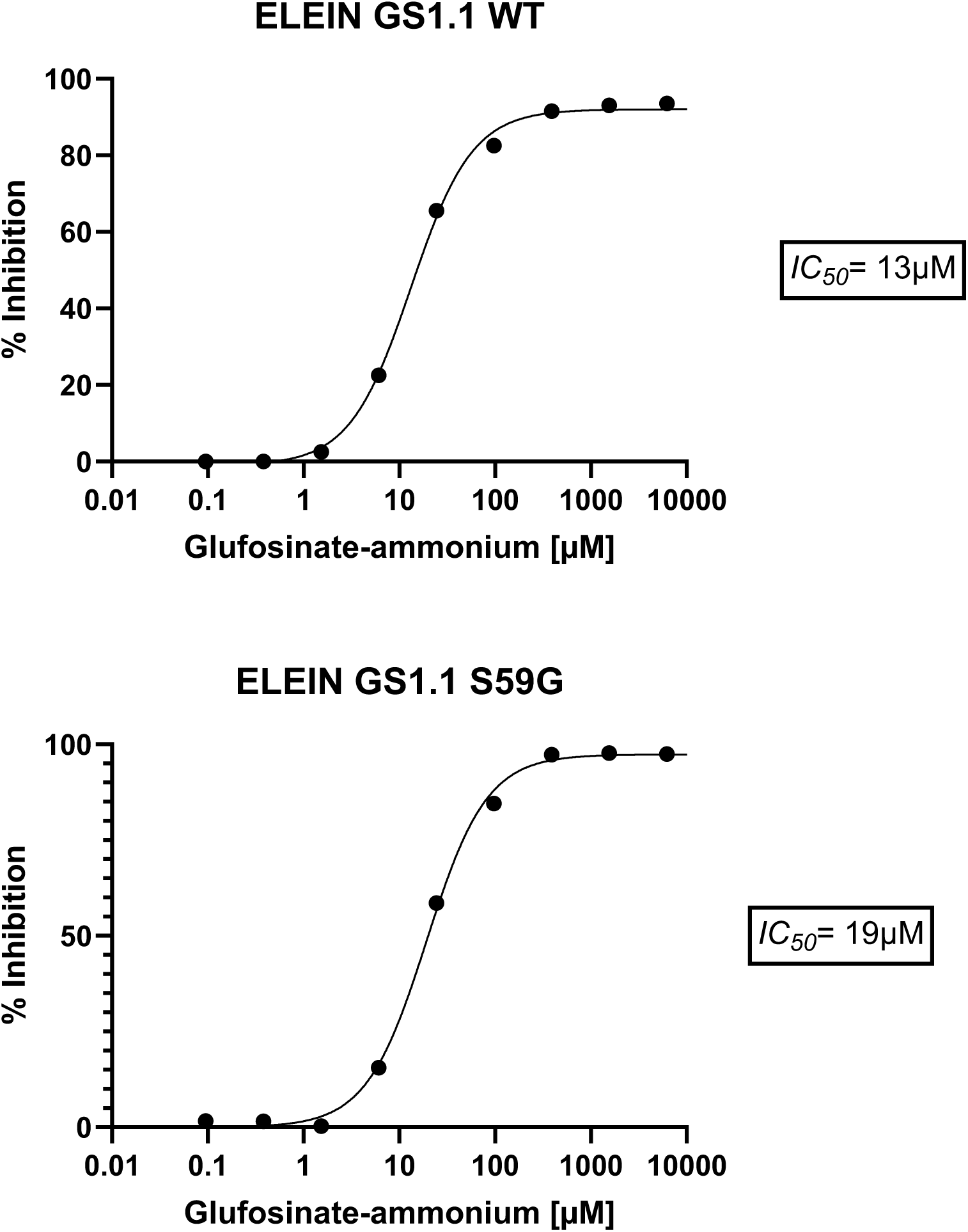
Inhibition of *Eleusine indica* GS1.1 wild-type (WT) and S59G mutant enzymes by glufosinate-ammonium (GFA). Enzyme inhibition was measured as the percentage decrease in activity relative to the uninhibited control across a range of GFA concentrations. Dose–response curves were fitted using a four-parameter logistic model. The WT enzyme (upper panel) exhibited an IC₅₀ of 13 μM, while the S59G mutant (lower panel) had an IC₅₀ of 13 μM. Data points represent the mean ± standard error of three replicate measurements.

**FIGURE 6.**
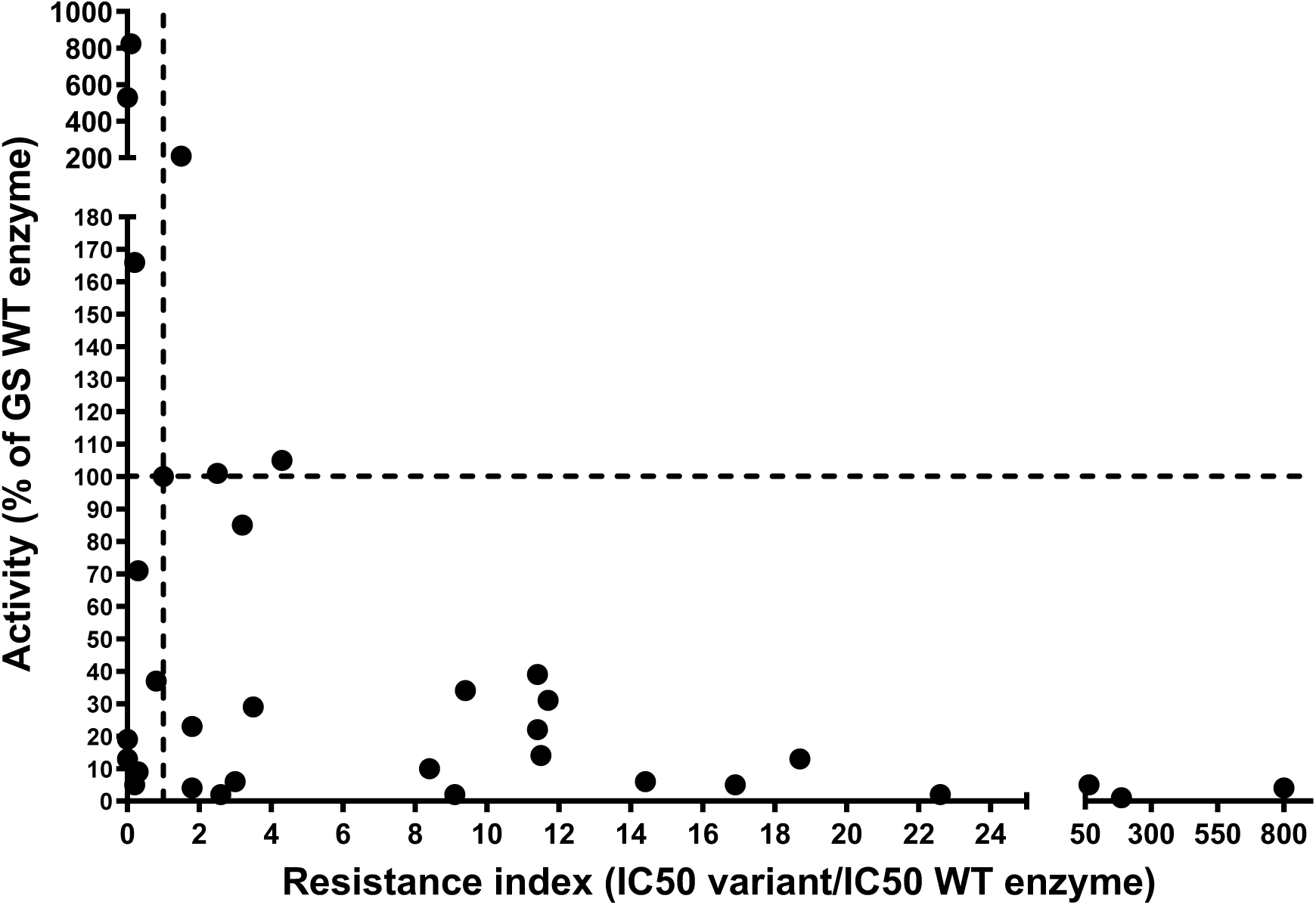
Relationship between resistance index (RI) and enzyme activity of *Amaranthus palmeri* chloroplastic GS1.1 variants.

## 4. Discussion

Digital PCR and sequencing of GS genes in the *A. palmeri* CCR population revealed a resistance profile distinct from that of previously characterized GFA-resistant populations such as MSR2.^28^ While the MSR2 population exhibited high-level amplification of GS2.1 and GS2.2 isoforms, consistent with robust overexpression of the GFA target site, the CCR population had minimal copy number variation. Most CCR individuals had GS copy numbers comparable to a susceptible reference (0.8–1.5-fold), with only a few plants, specifically CCR13, CCR17, and CCR22, displaying moderate increases in GS2.2 (up to ∼2.0-fold) and GS2.1 (Fig. 1A). GS CN variation was not prevalent among the assessed plants and did not exceed thresholds typically associated with gene amplification-driven resistance. This contrast shows the genetic variability of glufosinate resistance mechanisms across populations and highlights the CCR population’s unique profile lacking substantial GS amplification. There was no difference in GS expression between the resistant populations and the susceptible reference (Fig.1B).

The G255D substitution was identified in GS2.2 in heterozygous form in several CCR individuals. This residue is located adjacent to E251, a known contact site for GFA binding based on structural modeling.^29^ The substitution from glycine to aspartic acid introduces a charged residue in a conserved domain of the enzyme, which altered the active site conformation and reduced catalytic efficiency. There was a 42% decrease in GS activity for the G255D variant compared to wild-type GS2.2 (Fig. 2). In addition, GFA inhibition assays demonstrated that while the wild-type enzyme exhibited a typical dose-dependent inhibition curve (IC₅₀ = 1.1 × 10⁻⁴ M), the G255D variant showed no measurable inhibition across the tested concentrations, indicating complete insensitivity to GFA at the enzymatic level (Fig. 3).

However, when the G255D variant was ectopically expressed in *Arabidopsis thaliana*, it failed to confer GFA tolerance *in planta*. Transgenic lines expressing G255D exhibited similar injury symptoms and biomass reduction as wild-type plants across all GFA treatment rates (Fig. 4A). A parallel set of transgenics expressing the corresponding mutation in the Arabidopsis GS2.2 backbone (G254D) also showed no resistant phenotype (Fig. 4B).

Although G255D confers resistance to GFA at the enzyme level, it does not confer whole-plant resistance. This could be due to reduced enzyme efficiency, lack of compensatory mechanisms, or additional physiological constraints.

These observations emphasize the need for cautious interpretation of *in vitro* experiments when evaluating potential resistance-conferring mutations. The inability of G255D to confer GFA tolerance *in planta* implies its limited adaptive value when expressed in isolation. It remains to be determined whether this substitution represents a background polymorphism, an early-stage adaptation, or a mutation that requires co-occurring mechanisms to manifest a resistant phenotype. Overall, the resistance mechanism of the CCR population is unlikely to be based on target-site mutations or target overexpression.

Further insights into GS structure-function relationships can contextualize these findings. GS catalyzes the ATP-dependent assimilation of ammonia into glutamate to produce glutamine.^1,4^ Glufosinate acts as a competitive inhibitor of glutamate, binding at the same site due to structural similarity with the intermediate γ-glutamyl phosphate. Key residues involved in GFA binding, including E131, E192, G245, H249, R291, and R311, are conserved across GS isoforms.^29,30^ The G255D mutation lies in proximity to these residues and may influence local conformational dynamics affecting both substrate and inhibitor interaction.

To explore this further, different codon variants were introduced at key GS1.1 residues to evaluate their effects on GFA sensitivity and catalytic function. Inclusion of multiple substitutions at single positions like E131A/D/G/K, E192A/D/G/K/Q/V, G245N/R/V, H249L/N/Q/P/Y, R311G/L/P/Q, R322G/S/K/M/T/V (Table S3) allowed assessment of whether loss of herbicide sensitivity was a general outcome of amino acid replacement at these sites or specific to particular residue chemistries. The results indicate that although many of these variants resulted in loss of GFA sensitivity, they also reduced catalytic activity to negligible levels, which limits the potential of such substitutions to confer field-relevant resistance (Table 2). Additionally, variants obtained by molecular prediction method rendered similar results. Variants retaining more than 65% of wild-type catalytic activity and exhibiting a RI substantially higher than that of the GS2.2 G255D reference would warrant further evaluation for resistance; however, molecular docking analyses did not identify substitutions meeting these criteria (Fig. 6, Table S3). It is to be noted that while G255D was found in GS2.2, the panel of target site mutants was tested using the GS1.1 isoform backbone.

However, since GS1.1 and GS2.2 share a high sequence similarity, especially high conservation of the residues involved in GFA binding ^29,30^, it would be expected that target-site mutations tested in GS1.1 are likely to produce the same effect in GS2.2.

In a study involving soybean cell suspension culture, resistant lines displayed a 4.6-fold higher IC₅₀ of GS activity relative to untreated controls, indicating reduced sensitivity of the enzyme to inhibition. Sequencing revealed multiple nucleotide substitutions in the GS gene, including a H249Y change within the substrate/inhibitor binding region, consistent with altered target-site interaction.^25^ Although no enzyme stability assays were performed, the authors proposed that these substitutions may reduce enzyme stability while simultaneously diminishing herbicide-binding. Such mutations can be recovered under *in vitro* conditions, where minimal metabolic demands permit survival of impaired cells, but are unlikely to persist in field populations due to associated fitness costs. In contrast, DNA shuffling of rice GS1 produced a variant carrying the R295K substitution that conferred measurable GFA resistance when expressed in *Arabidopsis*.^26^ Although the level of resistance was weaker than that conferred by *bar/pat*, this study demonstrates the feasibility of engineering GS-based tolerance through specific amino acid substitutions. A study in *L. multiflorum* identified a D173N substitution in the GS2 gene that was proposed to confer GFA resistance.^31^ However, this hypothesis was disputed in a subsequent study reporting no difference in GS activity between resistant and susceptible biotypes.^23^ Structural modeling further demonstrated that residue 173 is distant from the catalytic site and unlikely to influence herbicide binding. All these studies illustrate the structural constraints of GS: some substitutions disrupt the herbicide binding but impair catalytic function, while others shift herbicide sensitivity without major loss of activity, emphasizing why GS mutations rarely translate into field-relevant resistance.

The GFA-resistant *E. indica* population originally described by Jalaludin et al. (2015)^10^ as exhibiting up to 20-fold resistance was subsequently investigated by Zhang et al. (2021).^24^ In that study, the S59G substitution in *EiGS1-1* was identified and functionally tested. However, the resistance conferred by this mutation was relatively small, with ∼1.5-fold shifts in crude extracts, ∼2-fold in yeast, and ∼2.5-fold in transgenic rice. This discrepancy indicates that S59G alone cannot account for the high resistance level reported by Jalaludin et al. As also acknowledged by Zhang et al., additional mechanisms are likely to contribute, potentially including enhanced antioxidant capacity or other non–target-site processes that mitigate GFA injury. Repeated assays in the present study did not reveal any IC₅₀ shift (Fig.5), in contrast to the 1.5- to 2.5-fold differences reported by Zhang et al.^24^ These results support the conclusion that the S59G substitution does not directly confer resistance. Such elevated activity could contribute to survival under sublethal doses of GFA, where inhibition of a proportion of GS enzymes may still allow sufficient residual activity in the more active mutant isoform.

Together, the apparent resistance in the mutant biotype arises not from reduced sensitivity but from increased catalytic activity of the mutant enzyme (Fig. S3) combined with efficacy limitations of GFA in *E. indica*.

Overall, this study constitutes a critical reference point for evaluating the functional consequences of GS mutations in herbicide resistance research. By demonstrating that G255D does not confer GFA resistance at the plant level, this work supports the ongoing effort to distinguish resistance-conferring mutations from non-functional polymorphisms in *A. palmeri* populations.

## 5. Conclusion

GFA resistance in the CCR *A. palmeri* population is not explained by GS amplification or the GS2.2 G255D substitution. While G255D abolished GFA inhibition *in vitro*, its reduced catalytic efficiency and failure to confer tolerance *in planta* highlight the strong functional constraints on target-site resistance. Broader mutational analyses confirmed that most substitutions disrupting GFA binding also compromise enzyme activity, underscoring the limited adaptive potential of GS alterations. These findings emphasize that GFA resistance in field populations is unlikely to arise from target-site modifications alone, reinforcing the need to investigate alternative, non–target-site mechanisms.

## Acknowledgements

The authors acknowledge BASF for funding and thank all co-authors for their contributions.

---

**Figure S1.**
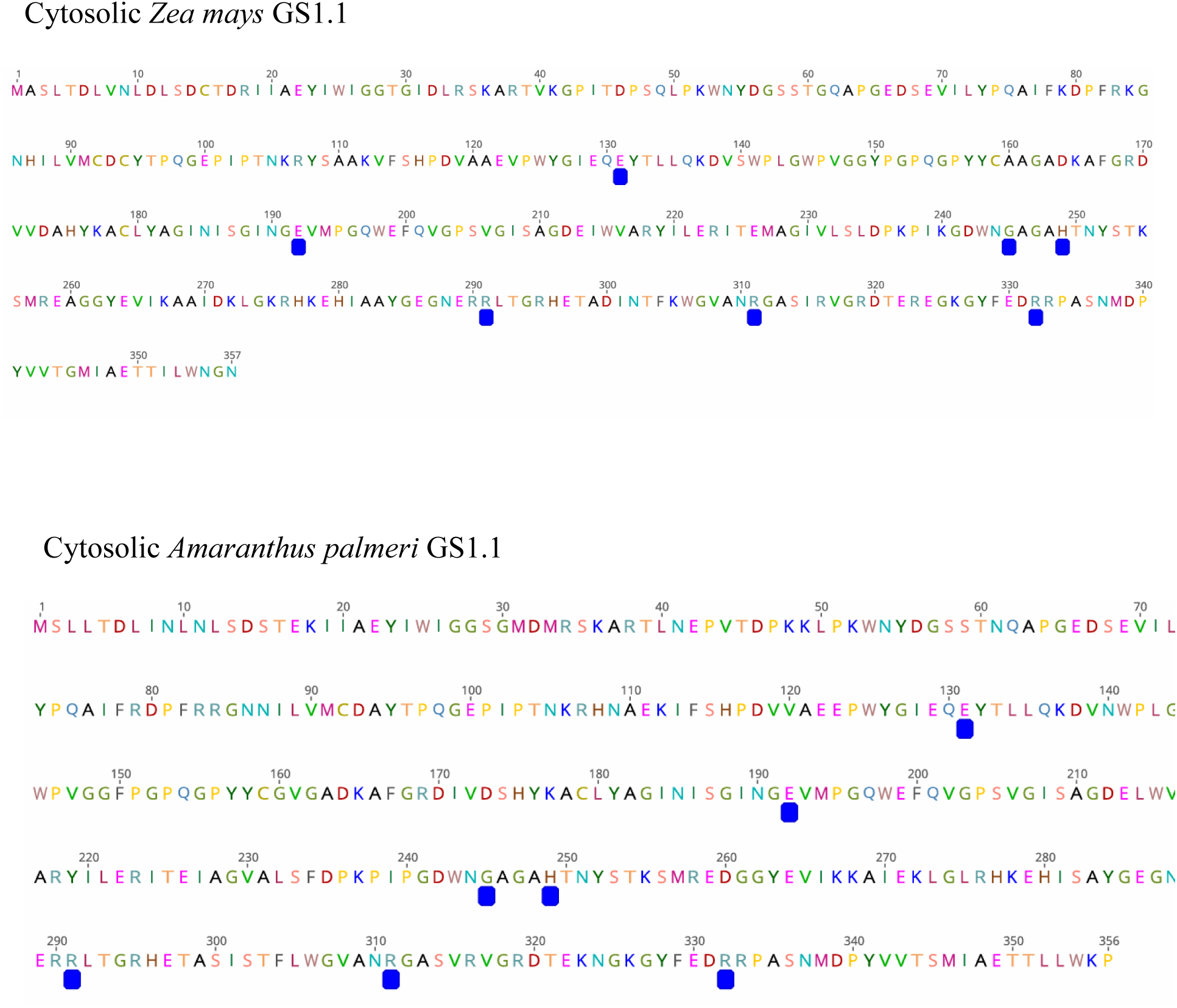
Glutamine synthetase 1 protein sequence of *Zea mays* (GS1.1) and *Amaranthus palmeri*. In blue squares are depicted the residues involved in P-PPT (phosphinothricin or glufosinate) as reported by Unno *et al*., 2006.

**Figure S2.**
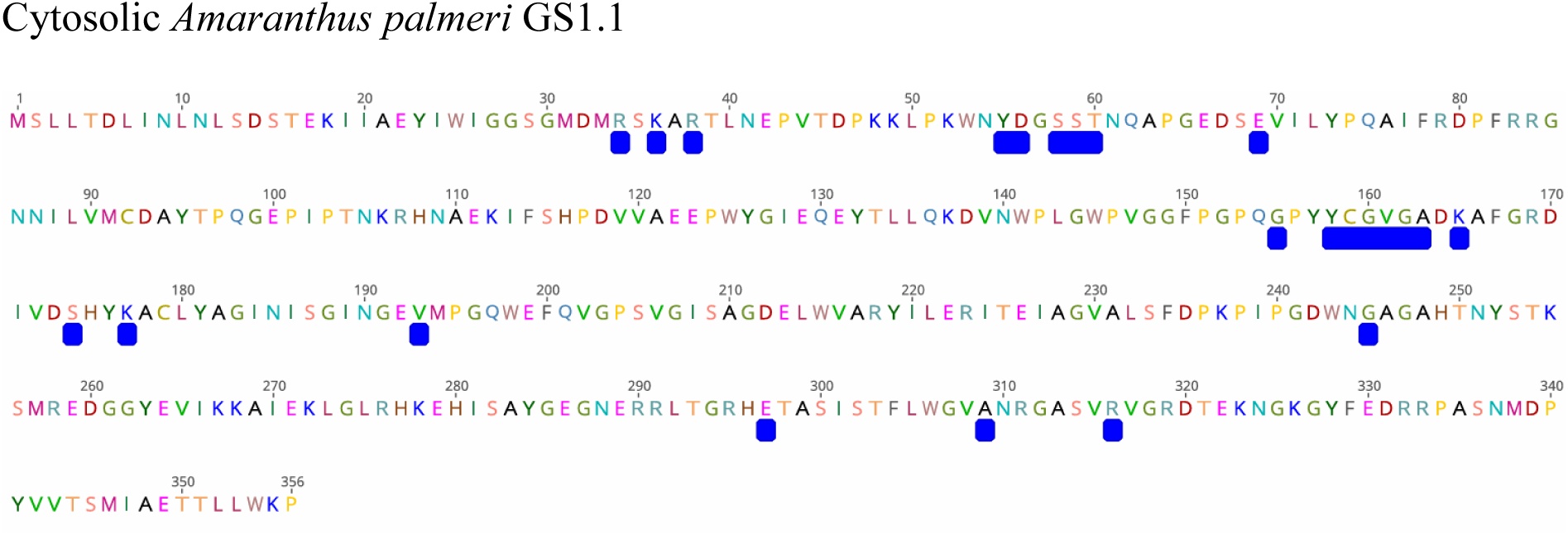
Glutamine synthetase 1 protein sequence of *Amaranthus palmeri*. In blue squares are depicted the residues involved in P-PPT (phosphinothricin or glufosinate) as obtained by docking simulation (see material and methods).

**Figure S3.**
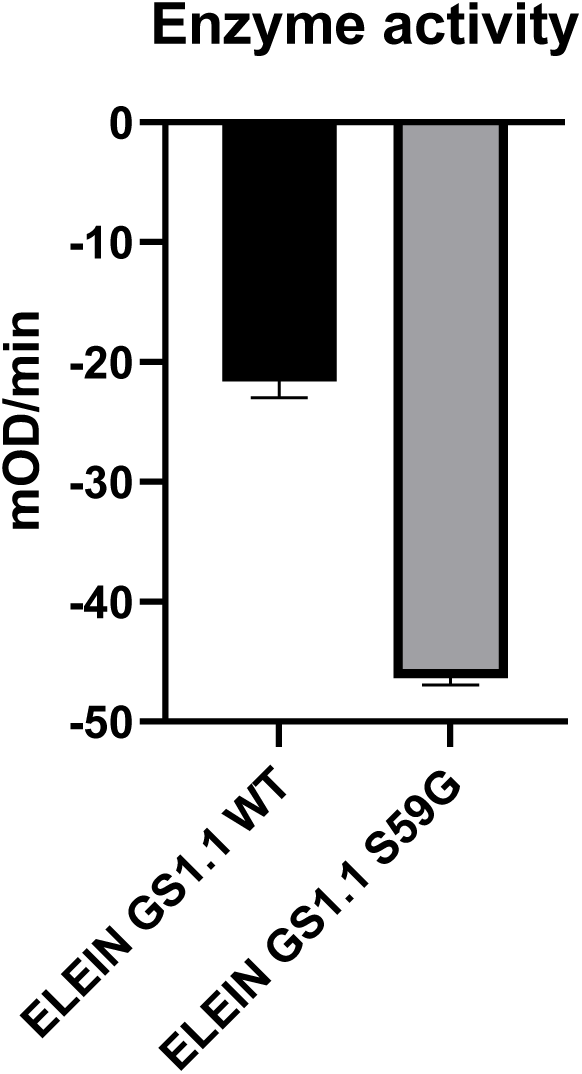
Enzyme activity of cytosolic glutamine synthetase 1 (GS1.1) from Eleusine indica. Activities of the wild-type (WT) and the S59G variant were determined by measuring the rate of absorbance change (mOD min⁻¹) in vitro. Bars represent mean values ± SE of three replicate assays.

**Table S1.**
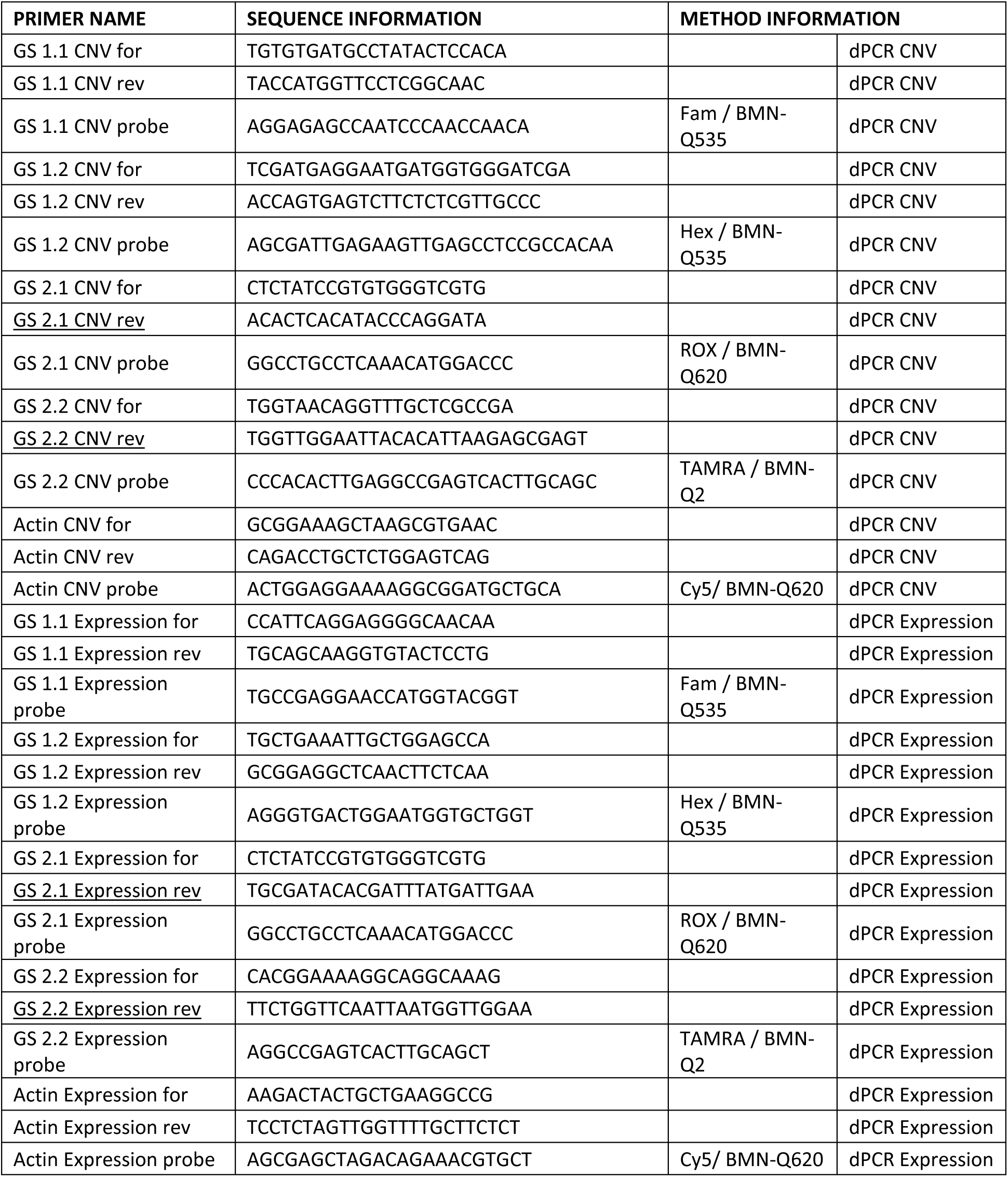
Primer information for copy number and expression analysis.

**Table S2.**
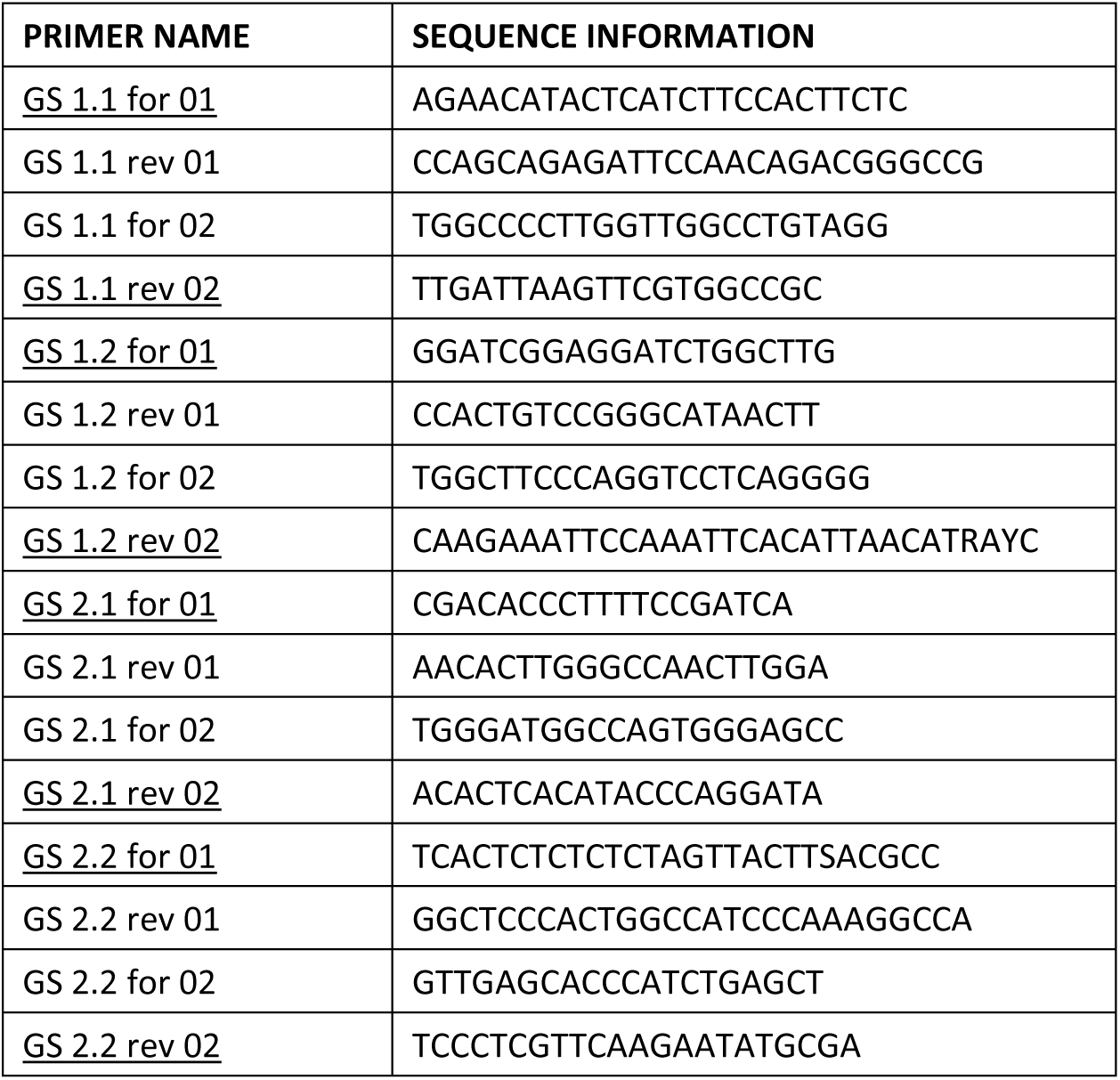
Primer information for cDNA sequencing.

**Table S3.**
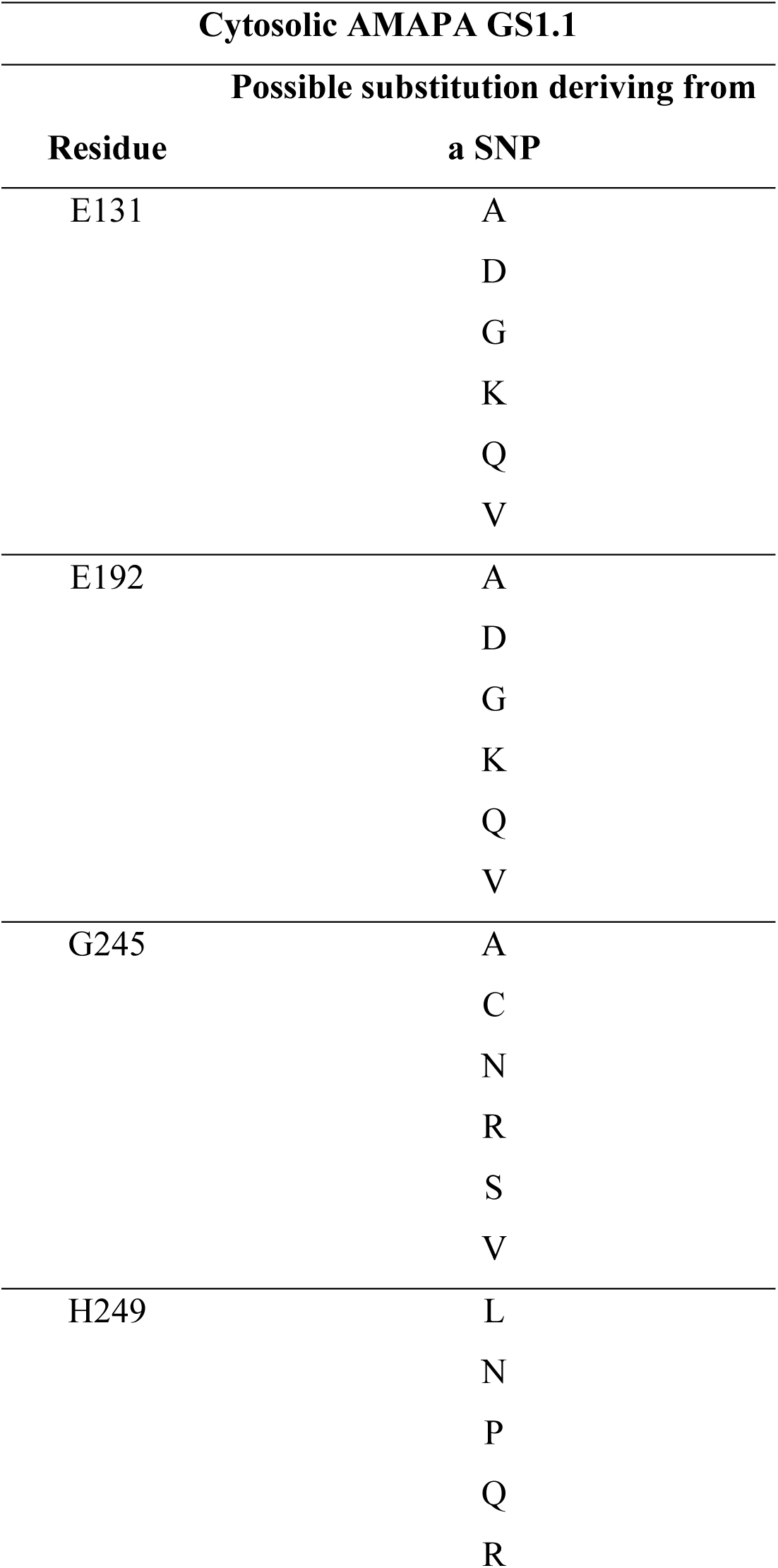
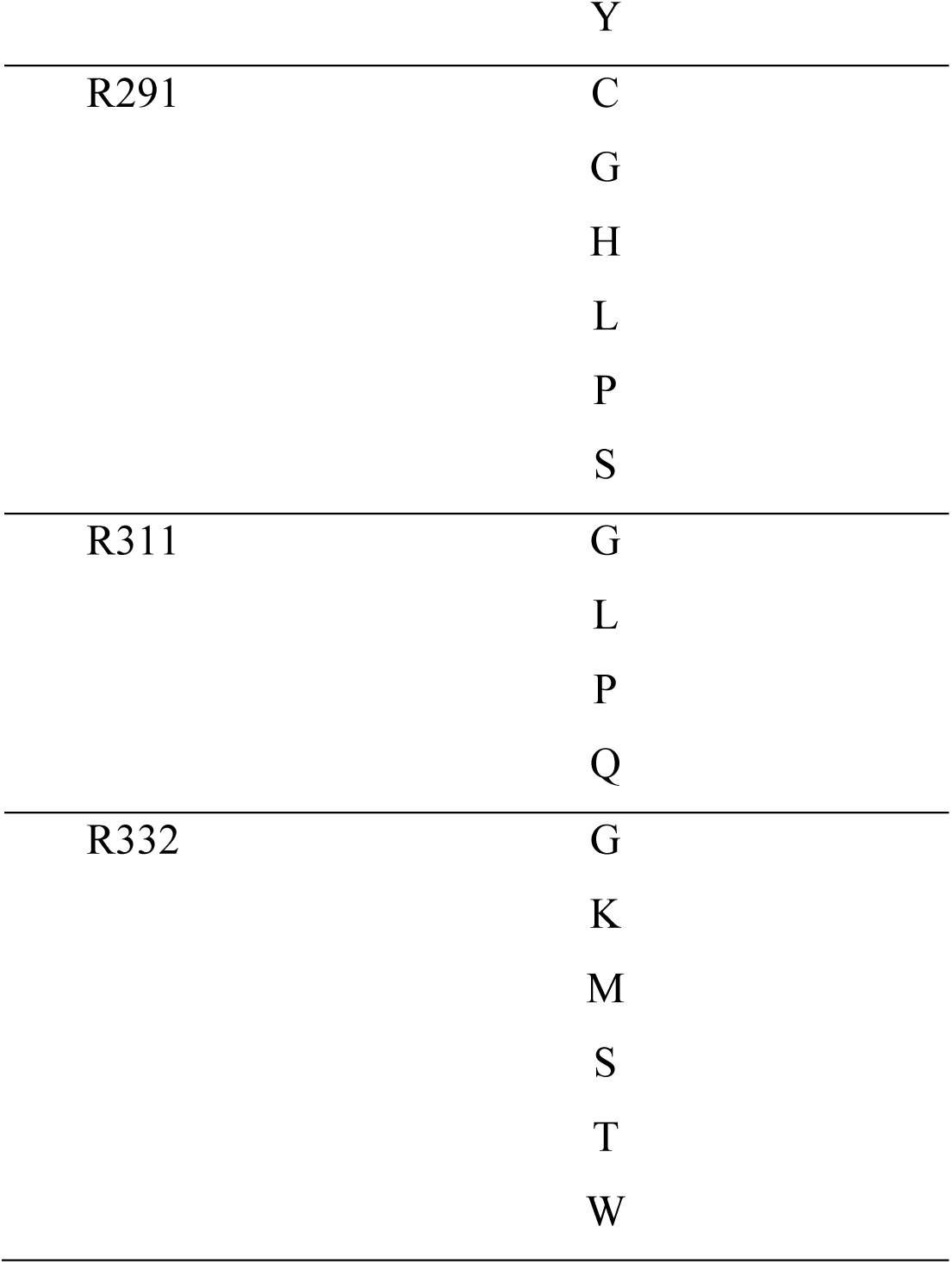
Possible amino acid substitutions in cytosolic Amaranthus palmeri GS1.1 resulting from single nucleotide polymorphisms (SNPs). The table lists residues in the GS1 protein known to interact with glufosinate (Unno *et al*., 2006). Each residue is followed by the set of alternative amino acids that could result from a SNP, based on one single base changes.

